# Recognition of familiar objects in tortoise hatchlings (*Testudo spp.*)

**DOI:** 10.1101/2021.03.22.436430

**Authors:** Silvia Damini, Gionata Stancher, Elisabetta Versace

## Abstract

Tortoises do not show parental care and live solitary except for the context of reproduction. Despite their limited need to interact with conspecifics, we previously observed that young tortoises, at their first experiences with conspecifics, can discriminate between familiar and unfamiliar conspecifics after just one encounter with another tortoise. Tortoise hatchlings ignored familiar conspecifics, while they first explored and then actively avoided unfamiliar conspecifics. It remains to be established whether the different reactions to unfamiliar and familiar individuals in tortoise hatchlings are reactions to novelty, or whether they are specific to the interactions with living animals. To test this, we familiarized one-month-old tortoise hatchlings with an object (a brown cone *vs*. a blue sphere) and then tested them in a novel arena once with the familiar object and once with an unfamiliar one. To measure the reactions toward familiar and unfamiliar objects, we measured the distance between the tortoise and the object throughout the test. Differently from what happened with unfamiliar and familiar conspecifics, we found no difference in behavior toward familiar and unfamiliar objects. This shows that the different reactions toward familiar and unfamiliar conspecifics previously observed are specific for social interactions and are not a mere reaction to the novelty effect. The behavioral responses displayed by young tortoises for unfamiliar conspecifics, but not for unfamiliar objects, show the relevance of social behavior from the beginning of life, even for solitary species.

## INTRODUCTION

Recognition is used by animals to discriminate among individuals and categorize them as offspring, mates, social partners, or neighbors (Yorzinski, 2017). Some animals can recognize individuals through familiarity by learning their specifics traits or by habituating to them after repeated interactions (Wiley, 2012). The ability to recognize familiar individuals can be useful in modulating future interactions with conspecific based on information gathered in previous encounters (Thom & Hurst, 2004). This ability is documented in social animals of different taxa, such as fish (Miklosi et al., 1997), mammals (Brennan & Kendrick, 2006), birds (Beer, 1971), and invertebrates (Tibbetts, 2002). There is also evidence of the ability to recognize familiar conspecifics in adult reptiles, particularly in the contest of territoriality, through visual and olfactory cues (Whiting, 1999; Carazo et al., 2008).

Evidence of the ability to discriminate between familiar and unfamiliar individuals at the beginning of life in non-social species has been recently found in tortoises’ hatchlings (Versace et al., 2018). Finding this ability in young and inexperienced tortoises has given insights into the ecological value and the evolutionary history of this social skills. For this reason, we want to further investigate the ability to recognize familiar individuals in young tortoises.

Tortoises are a clade of terrestrial turtles whose evolutionary relationships remain to these days poorly known (Parham et al., 2006) but we know that for the past 30 million years they evolved without parental care (Joyce, 2013). Other indications of their solitary nature are the fact that females increase their mobility just before laying eggs to disperse nests on a wider area (Diaz-Paniagua et al., 1996) and the ranges of hatchlings have little overlap (Keller et al., 1997). Moreover, tortoises mate promiscuously and do not form pair bonds or cohesive social groups (Ernst & Barbour, 1989; Pearse & Avise, 2001). In wild tortoises, evidence of social interactions is mostly limited to behaviors that are performed when sexual maturity is reached, years after hatching, such as courtship, mounting, and nesting (Auffenberg, 1977; Galeotti et al., 2005; Sacchi et al., 2003).

Despite these solitary habits, mounting evidence suggests that tortoise hatchlings are sensitive to social stimuli from the beginning of life (Versace et al., 2020). Versace et al. (2018) studied the behavior of tortoise hatchlings with limited social experience when exposed to familiar and unfamiliar conspecifics. We observed that hatchlings previously exposed to just one conspecific behave differently toward familiar or unfamiliar conspecifics. Looking at the distance between tortoises, we observed that tortoises ignored familiar conspecifics (they progressively approached the distance expected by random trajectories) while in front of unfamiliar conspecifics they first explored and then actively avoided them. These differences in behavior were interpreted as evidence of discrimination between familiar and unfamiliar individuals. However, it remains to be clarified whether tortoise hatchling exhibited reactions specific to social partners or whether they simply reacted to the novelty of unfamiliar objects. Species like rats and mice tend to interact more with novel objects than with familiar ones (Bevins & Besheer, 2006), this behavior is known as the novelty effect. This effect is used in the novel object recognition test to investigate various aspects of cognition in animals. It can be used to study memory and learning, the preference for novelty, the influence of different brain regions in the process of recognition, and even the study of different drugs and their effects (Antunes & Biala, 2012).

To clarify whether the reaction to unfamiliar conspecifics observed in tortoise hatchlings is present only in the context of interaction with other animals or if it is present also in the interaction with inanimate objects, we performed an experiment using the same method used by Versace et al. (2018) but using familiar and unfamiliar objects as experimental manipulation. We familiarized and tested tortoise hatchlings of four species (*Testudo marginata, Testudo graeca*, a Hybrid between these two, *Testudo hermanni*, and *Testudo horsfieldii*) with objects instead of conspecifics. After hatching in individual compartments, tortoises were raised in separate boxes together with an object (either a brown cone or a sphere), before testing them in a novel environment together with the familiar object (familiar condition) or unfamiliar one (stranger condition). Most chelonians have good vision thanks to their complex eyes that allow them to distinguish images clearly and perceive colors (Granda & Stirling, 1965). Herman’s tortoises can discriminate between colors and have a preference for yellow, red, and violet stimuli (Pellitteri-Rosa et al., 2010). We avoided these colors because they could have added confounding variables to the experiment.

If the different behavior displayed by tortoise hatchings with familiar and unfamiliar individuals was just a consequence of the novelty of the object, we expected to see a difference also between the familiar object and the unfamiliar object conditions. Instead, if the differences observed with unfamiliar and familiar individuals are only present in the context of social interactions, we expected to observe no difference between the familiar and unfamiliar objects.

## METHODS

### Subjects and rearing conditions

We observed 37 newly hatched tortoises of four species: 7 *Testudo hermanni*, 6 *Testudo horsfieldii*, 12 *Testudo graeca*, 4 *Testudo marginata*, and 8 hybrids between *Testudo graeca* and *Testudo marginata* (Hybrid from now on). Tortoises were about one month old at the moment of the test (24-56 days of age, average 38 days). Eggs laid outdoor on the ground by tortoises were collected by the experimenters and incubated in darkness at 31 °C +/-2 °C. Tortoises hatched in darkness in individual compartments and then were paired with one of the two experimental objects (1-41 days of age, average 19 days). Tortoises were fed with green leaves and hydrated at least twice daily. After hatching, each subject was housed in a square-shaped arena (for the subjects 1-2-3 the size was 15×15×12 cm, for the others 20×20×12 cm) with the bottom covered with soil, leaves, and straw. The object was placed in the center of the arena, so that could be easily inspected on all sides. Before the test, subjects had never seen the unfamiliar object.

### Stimuli

We used two wood objects as familiarization and test stimuli: a light blue sphere of 3.5 cm in diameter and a brown cone of 4.5 cm in height (Fig. 1). Each object was glued to a square support of 3.5 cm of side that was buried in the soil to maintain the object in place.

**Figure 1.**
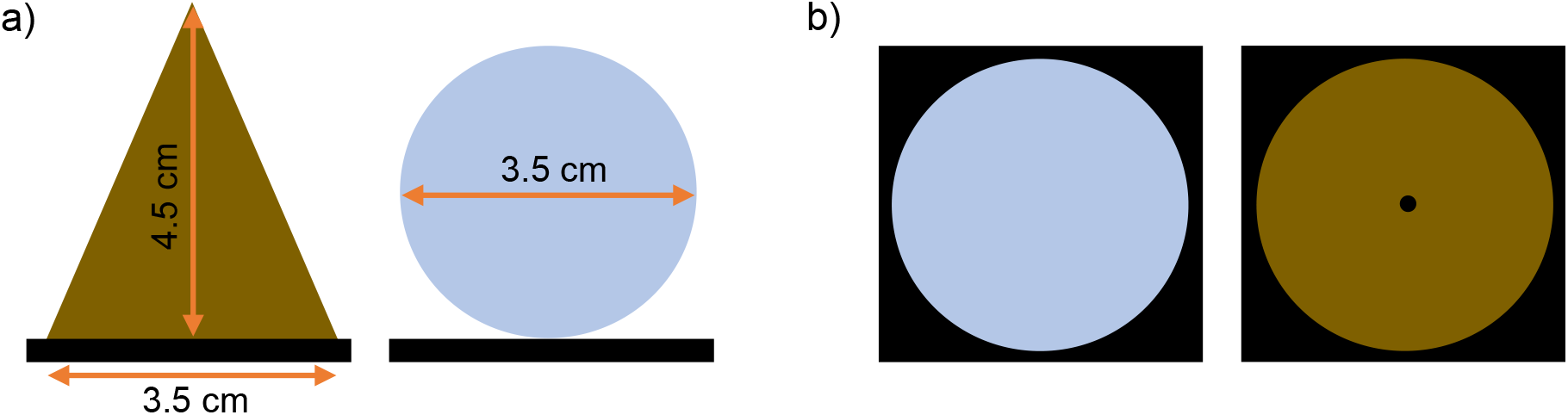
a) Schematic representation of the experimental objects on their support from the side and measurement. b) Schematic representation of the experimental objects on their support from above.

### Experimental apparatus

The experimental apparatus was similar to the one used in Versace et al. 2018 (Fig. 2a). We used a circular arena (∅ 25 cm, 10 cm high, Figure 2a) with the bottom covered with wet sand (0.5 cm). A webcam located on top of the center of the arena recorded tortoises’ behavior.

**Figure 2.**
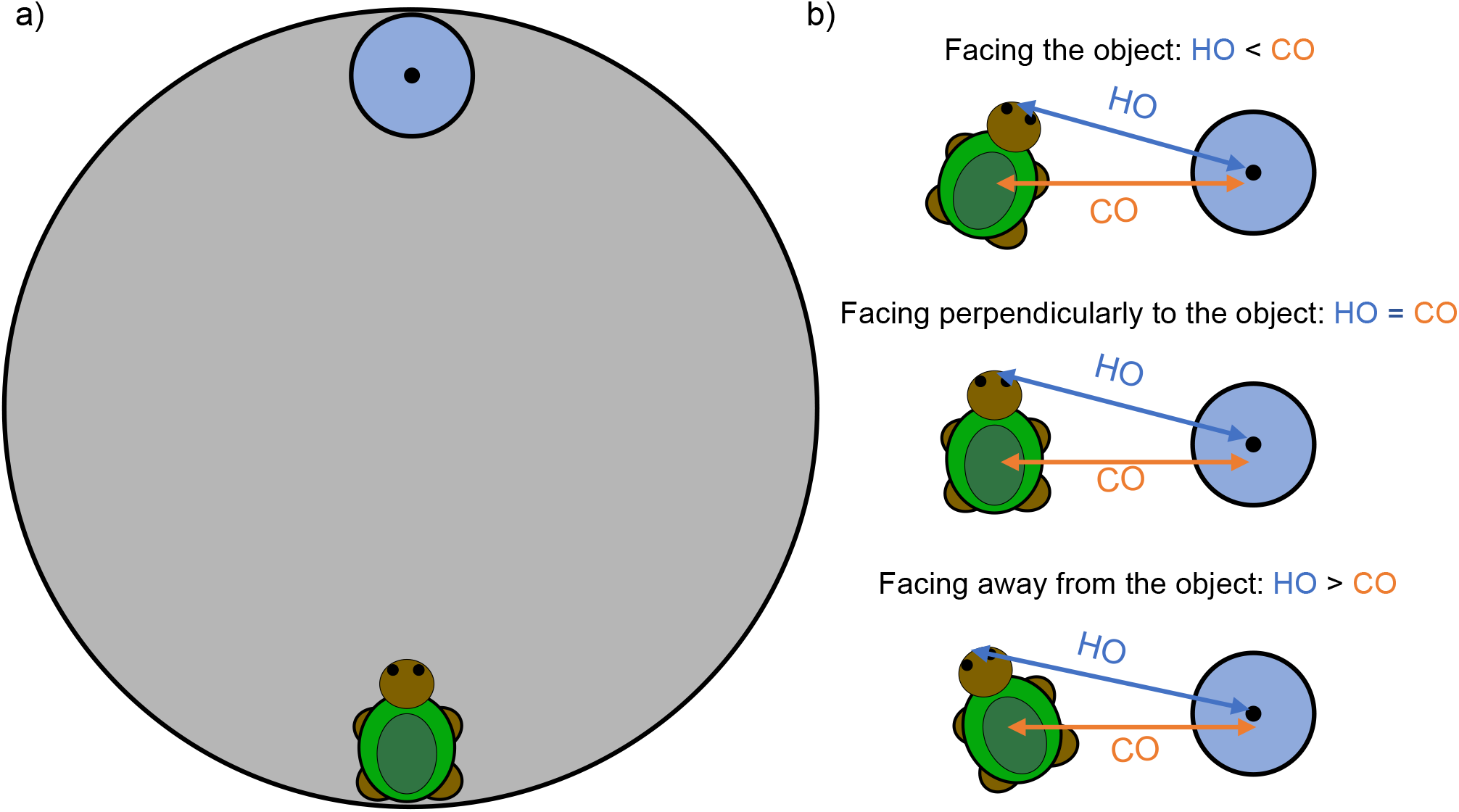
a) Schematic representation of the experimental apparatus with a subject and one of the objects located in the starting position. b) Distances between the head (H)/carapace (C) and object (O). When HO<CO the subject is facing the object when HO=CO the direction of the subject is perpendicular to the object when HO>CO the subject is facing away from the object.

### Procedure

The procedure was divided into two phases: familiarization and test. Both phases were similar in duration and modalities to the ones described in Versace et al. (2018). In the familiarization phase, we raise tortoises in an enclosure with one object placed in the center. Each subject remained with the object for at least 14 days (14-32 days, average 19.5 days) before the test, but the time varied depending on experimental needs and atmospheric conditions (tortoises wouldn’t be active enough to be tested on cloudy days).

In the test phase, we counterbalanced the order of test with the familiar/unfamiliar object (19 tortoises were tested first with the familiar object the other 18 tortoises were tested first with the unfamiliar object). At least 2 days elapsed between the two trials (2-14 days, average 6 days). Each tortoise was tested once with each stimulus (except subject 20 which was tested only for the familiar condition). The list of tests is shown in Table 1.

**Table 1.**
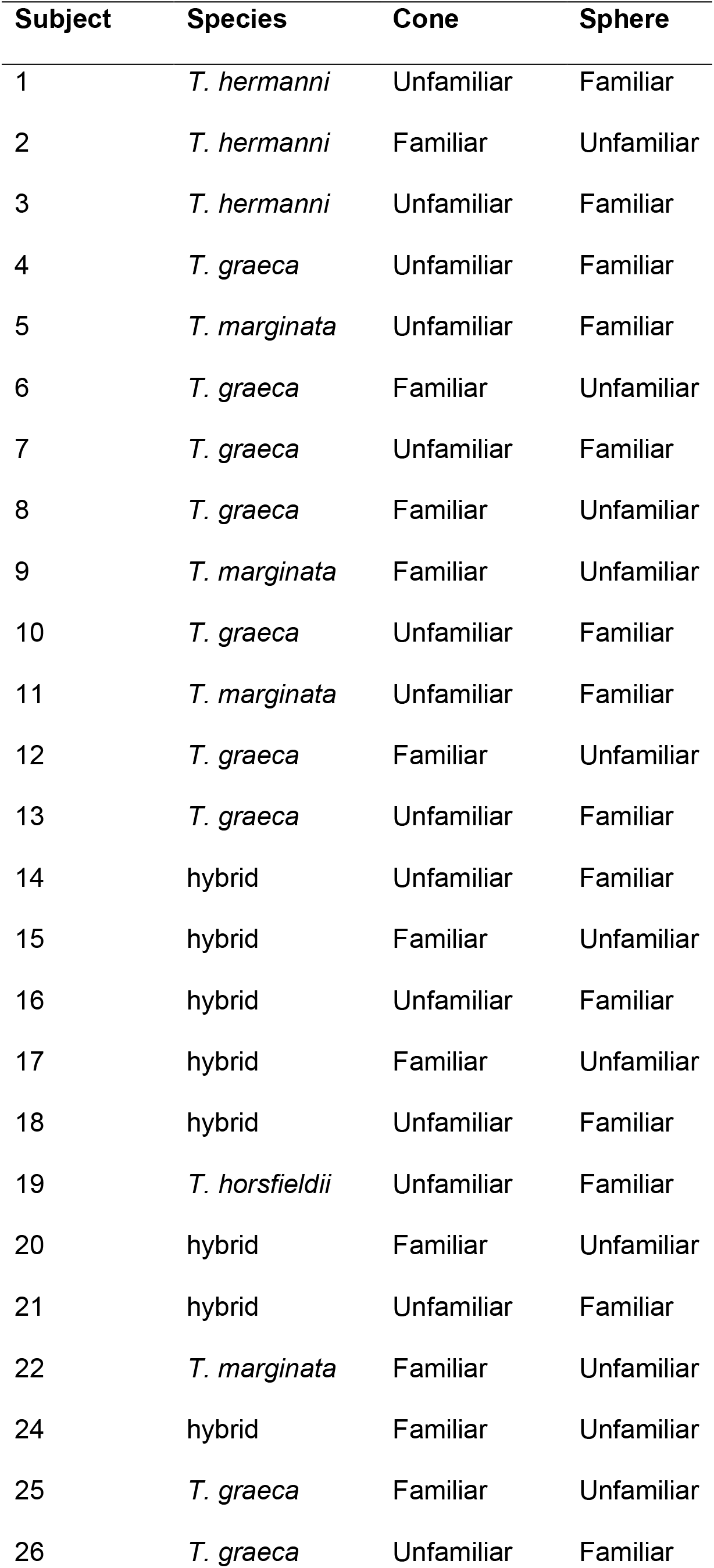
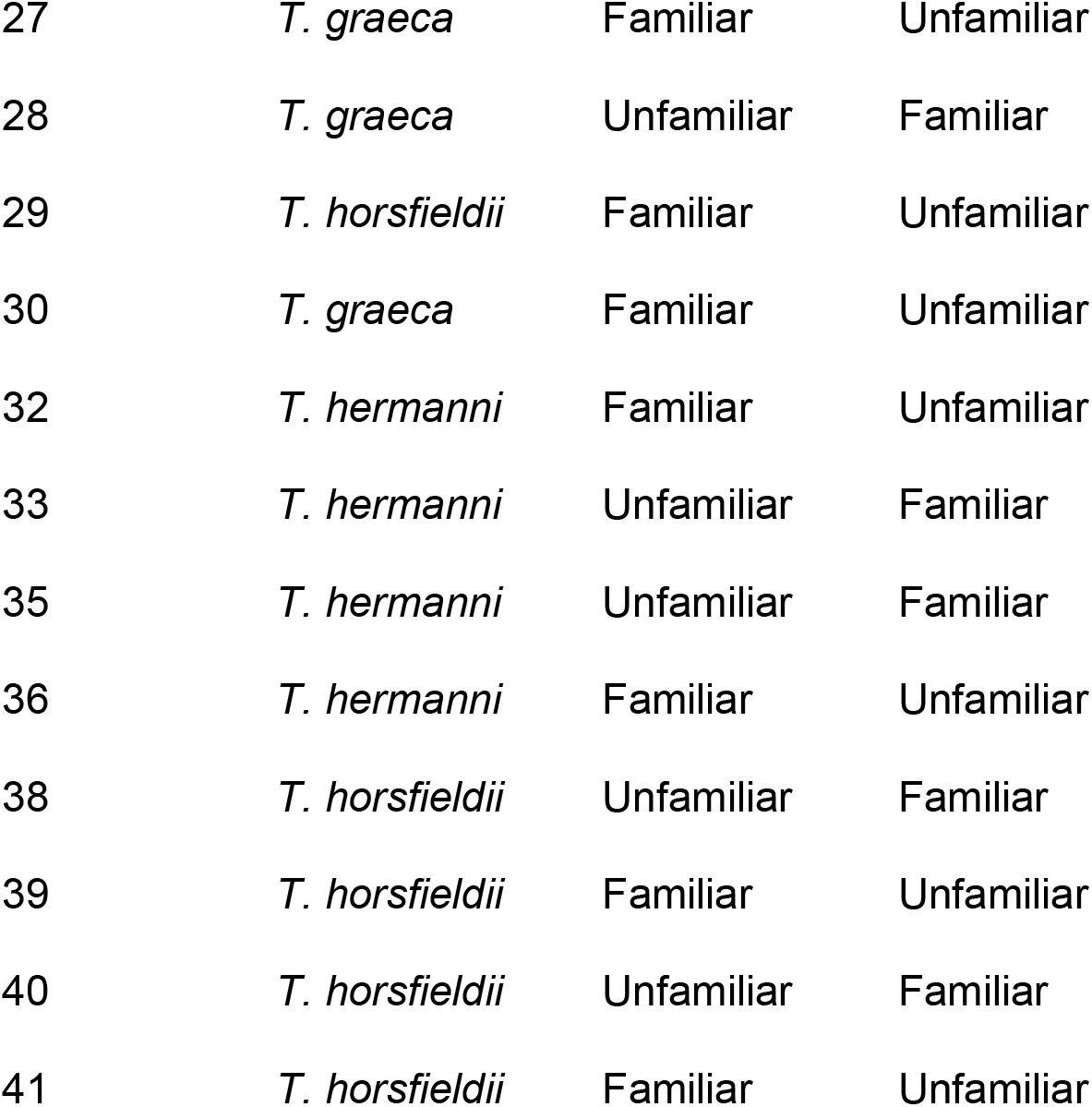
List of experimental subjects by Species and Condition (referred to the object).

| Subject | Species | Cone | Sphere |
| --- | --- | --- | --- |
| 1 | *T. hermanni* | Unfamiliar | Familiar |
| 2 | *T. hermanni* | Familiar | Unfamiliar |
| 3 | *T. hermanni* | Unfamiliar | Familiar |
| 4 | *T. graeca* | Unfamiliar | Familiar |
| 5 | *T. marginata* | Unfamiliar | Familiar |
| 6 | *T. graeca* | Familiar | Unfamiliar |
| 7 | *T. graeca* | Unfamiliar | Familiar |
| 8 | *T. graeca* | Familiar | Unfamiliar |
| 9 | *T. marginata* | Familiar | Unfamiliar |
| 10 | *T. graeca* | Unfamiliar | Familiar |
| 11 | *T. marginata* | Unfamiliar | Familiar |
| 12 | *T. graeca* | Familiar | Unfamiliar |
| 13 | *T. graeca* | Unfamiliar | Familiar |
| 14 | hybrid | Unfamiliar | Familiar |
| 15 | hybrid | Familiar | Unfamiliar |
| 16 | hybrid | Unfamiliar | Familiar |
| 17 | hybrid | Familiar | Unfamiliar |
| 18 | hybrid | Unfamiliar | Familiar |
| 19 | *T. horsfieldii* | Unfamiliar | Familiar |
| 20 | hybrid | Familiar | Unfamiliar |
| 21 | hybrid | Unfamiliar | Familiar |
| 22 | *T. marginata* | Familiar | Unfamiliar |
| 24 | hybrid | Familiar | Unfamiliar |
| 25 | *T. graeca* | Familiar | Unfamiliar |
| 26 | *T. graeca* | Unfamiliar | Familiar |
| 27 | *T. graeca* | Familiar | Unfamiliar |
| 28 | *T. graeca* | Unfamiliar | Familiar |
| 29 | *T. horsfieldii* | Familiar | Unfamiliar |
| 30 | *T. graeca* | Familiar | Unfamiliar |
| 32 | *T. hermanni* | Familiar | Unfamiliar |
| 33 | *T. hermanni* | Unfamiliar | Familiar |
| 35 | *T. hermanni* | Unfamiliar | Familiar |
| 36 | *T. hermanni* | Familiar | Unfamiliar |
| 38 | *T. horsfieldii* | Unfamiliar | Familiar |
| 39 | *T. horsfieldii* | Familiar | Unfamiliar |
| 40 | *T. horsfieldii* | Unfamiliar | Familiar |
| 41 | *T. horsfieldii* | Familiar | Unfamiliar |

Before the beginning of the experimental session, we regulated the external temperature of the subject under a light, to make sure it was active. Subsequently, the experimental subject was located at a diametrically opposed position from the object (familiar/unfamiliar, according to the experimental condition), facing the center of the arena (Fig. 2a). The behavior was recorded for 15 minutes from the moment in which the tested tortoise moved the first step (a movement of at least one leg that displaced the carapace). If the tortoise did not move for more than 10 minutes, the session was aborted and repeated the subsequent day.

### Data analysis

To score the behavior, we extracted one frame every 20 seconds (3 frames per minute) and used ImageJ (Rasband, 2017) to identify the location of the center of the carapace, the tip of the head of the tortoise, and the center of the object in all frames.

For every test, we calculated, for 5 consecutive periods of three minutes (15 minutes overall), the distance between the centroids of the carapaces (C) and the object (O) in centimeters (CO), the HO distance between the tip of the heads (H) and the object (O) in centimeters (HO), and the difference between these measures (CO-HO) (Fig. 2b).

The tortoise-object distance provided a measure of proximity irrespective of the relative orientation of the tortoise. The difference between the distance of the centroids from the object and the distance of the heads from the object provided an index of the relative orientation of the subject: if the distance between the head of the subject and the object is bigger than the distance between the carapace of the subject and the object the tortoise is facing away from the object if the distance between the head of the subject and the object is smaller than the distance between the carapace of the subject and the object the tortoise is facing the object (Fig. 2b).

We analyzed variation of the Distance between tortoise and object, looking at the effect of the independent variables Condition (familiar vs. unfamiliar), Species (*T. hermanni, T. horsfieldii, T. graeca, T. marginata*, and *Hybrid*), and Time (0,1-3, 4-6, 7-9, 10-12, 13-15 minute) as fixed effects and Subject as random effect To analyze the facing orientation we created an orientation coefficient dividing the difference between the distance between the carapace of the tortoise and the object and the distance between the head of the tortoise and the object for the sum of the distance between the carapace of the tortoise and the object and the distance between the head of the tortoise and the object. Negative values indicate facing away from the object, positive values indicate facing the object.

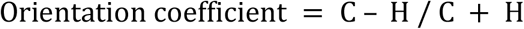

The facing orientation was analyzed using a mixed-design regression with Condition, Species, and Time as fixed effects and Subject as random effect. Analyses were conducted with R (version 3.2.1) using the libraries ggplot2, nlme, ez, car Rmisc, and multcomp.

### Ethical Note

The experimental procedures were approved by the Ethical Committee of the Fondazione Museo Civico Rovereto (Italy). Experiments comply with the current Italian and European Union laws for the ethical treatment of animals and are in line with the ASAB/ABS Guidelines for the Use of Animals in Research.

## RESULTS

### Tortoise-object distance and facing orientation

The tortoise-object distance was analyzed performing a mixed design regression including Time, Species, and Condition as independent variables and Subject as random effect. There was a significant main effect of Time (F(5,155) = 63.051, p < 0.001) and of interaction between Species x Time (F(20,155) = 1,855, p < 0.05) and Species x Condition (F(4,186) = 6.264, p < 0.001). There was no significant main effect of Condition (F(1,186) = 0.246, p < 0.621) and Species (F(4,31) = 0.615, p = 0.655), nor of interaction between Condition x Time (F(5,186) = 0.362, p = 0.874) and interaction between Species x Condition x Time (F(20,186) = 1.192, p < 0.265). In both the Familiar and the Unfamiliar conditions tortoises progressively approached the object fast at the beginning of the test and then maintained approximately the same distance toward the end (see Fig. 3).

**Figure 3.**
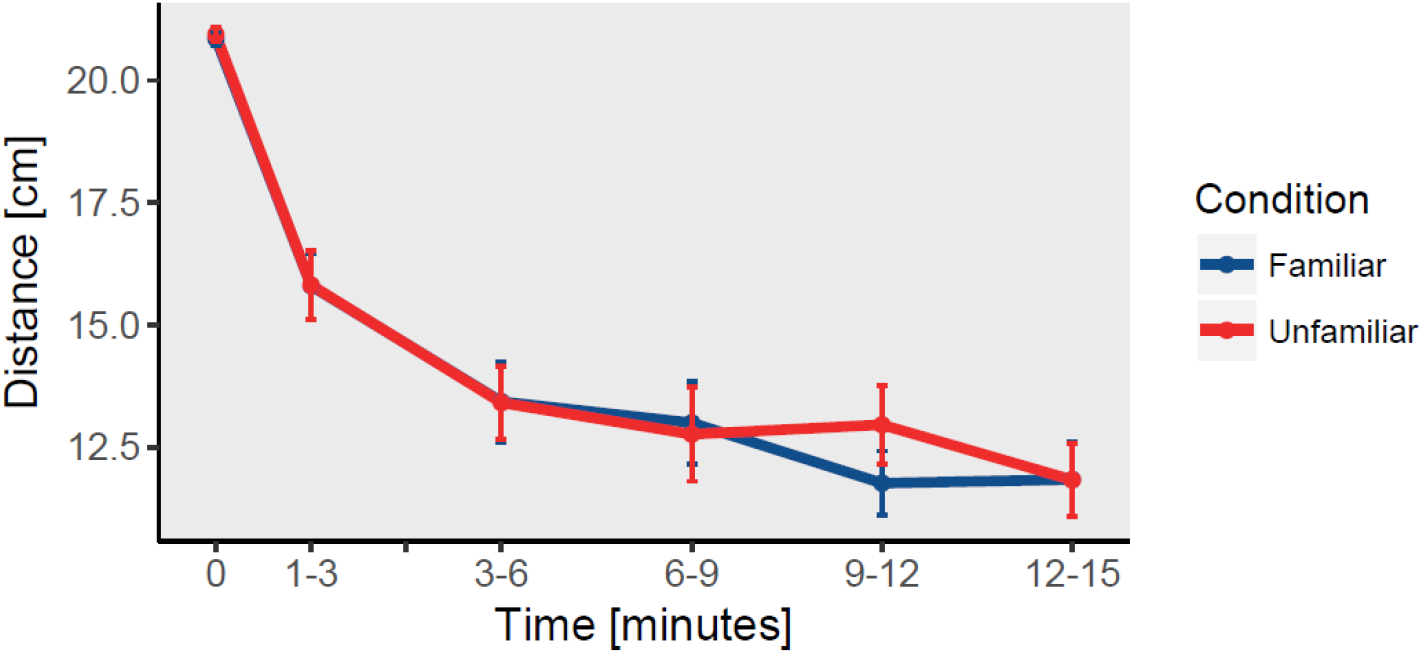
Average carapace distance between tortoises’ centroids and object in centimeters by Condition in Time. At the beginning of the experiment (Time 0) the subject was placed diametrically opposed to the object, then the tortoise-object distance was recorded at five time points (1-3, 3-6, 6-9, 9-12, 12-15). Blue indicated the familiar object condition while red indicated the unfamiliar object condition.

There was a significant main effect of Time (F(5,155) = 9.445, p < 0.001). There was no significant main effect of Condition (F(1,186) = 1.624, p = 0.204) and Species (F(4,31) = 0.572, p = 0.685), nor for interaction between Species x Time (F(20,155) = 0.888, p = 0.603), Species x Condition (F(4,186) = 0.274, p = 0.895), Condition x Time (F(5,186) = 0.293, p = 0.916), Species x Condition x Time (F(20,186) = 0.166, p = 1). In both the Familiar and the Unfamiliar conditions tortoises looked less toward the object as the test progressed. See graph in Fig. 4.

**Figure 4.**
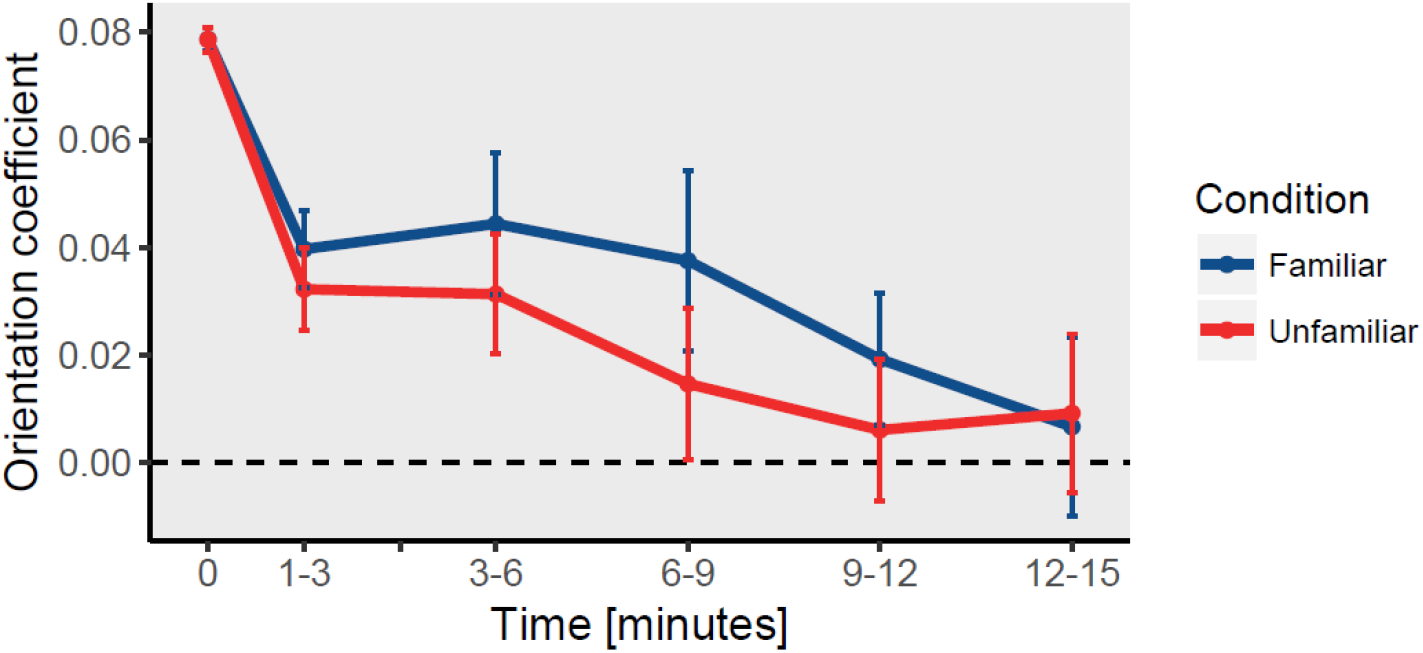
Facing orientation of the tortoise to the object by Condition in Time. At the beginning of the experiment (Time 0), the subject was placed facing toward the object, then the tortoise-object orientation coefficient was calculated at five time points (1-3, 3-6, 6-9, 9-12, 12-15). Positive values indicate that the tortoise is facing the object, negative values indicate that the tortoise is facing away from the object. Blue indicated the familiar object condition while red indicated the unfamiliar object condition.

## DISCUSSION

This experiment aimed to verify whether the fact that tortoise hatchlings react more to unfamiliar conspecifics while ignoring familiar ones (Versace et al., 2018) is a consequence of the novelty of the stimuli (Berlyne, 1950) or if it is a behavior only present in the contest of social interactions.

We observed that tortoise hatchlings do not behave differently in the presence of familiar or unfamiliar inanimate objects. Both in the presence of familiar and unfamiliar objects, hatchling got closer to the object over time. This happened particularly quickly at the beginning of the test. Moreover, the subjects oriented their attention to the object as the test progressed in time. These behaviors did not differ in the familiar and unfamiliar conditions. Differently from when hatchlings had been tested with conspecifics, we observed a consistent interest to face the objects and non-significant trend indicating a potential greater interest for familiar objects (hence in the opposite direction of what has been observed for hatchling located in front of familiar/unfamiliar conspecifics). This excludes the possibility that the differences in behavior toward familiar and unfamiliar conspecifics can be explained by the novelty effect. These findings are in line with the hypothesis that tortoise’s tendency to avoid unfamiliar conspecifics is specific to the context of interaction with living beings.

Other pieces of evidence of social abilities in tortoises are their spontaneous preference for face-like stimuli (Versace et al., 2020), and their gaze following (Wilkinson et al., 2010), as well as social learning abilities (Wilkinson et al., 2010). All these findings are in line with the idea that even non-social animals possess adaptations that allow them to interact with and learn from other animals present in their environment. Some of these traits are present at the beginning of life, pointing at the early onset of social skills in tortoise hatchlings, despite their solitary life. Other experiments on the recognition of familiar individuals in animals without extended social experience have been performed on chicks through filial imprinting (Vallortigara & Andrew, 1994; Zajonc et al., 1975). Differently from tortoises, domestic chicks are social animals, that live in flocks and rely on parental care (Nicol, 2015). Our results suggest that the ability to discriminate between familiar and unfamiliar individuals might have evolved in contexts different than repeated social interactions and that even in non-social species selection for social/asocial behavior is at work at the onset of life.

## Authors’ contributions

E.V., S.D., and G.S. conceived the project; E.V., S.D., and G.S. designed the experiment; S.D. carried out the experiment; S.D. analyzed the data; S.D. drafted the paper; all authors revised the manuscript and gave final approval for publication.

## Acknowledgments

We thank the internship students of the high school “Liceo Antonio Rosmini” (Rovereto, Italy) for helping in data collection and the Rovereto Civic Museum Foundation, for providing the facilities to carry out the present research.

